# Laminin and BDNF synergistically induce local translation in axonal growth cones

**DOI:** 10.64898/2026.02.09.704908

**Authors:** Nikita Kirkise, Kristy Welshhans

## Abstract

The laminins are a family of extracellular matrix proteins that regulate numerous cellular processes, including adhesion, neurite outgrowth, and axon guidance. However, it remains unclear whether laminin regulates axon guidance through local translation. Here, we show that laminin is necessary for local translation in axonal growth cones. Local translation is significantly increased in growth cones of embryonic day 17 mouse cortical neurons, either cultured on or acutely stimulated with soluble laminin 111, in the presence of BDNF. When cultured on laminin isoforms 211 or 221 in the presence of BDNF, there was a remarkable decrease in local translation in growth cones. Using a puromycin-proximity ligation assay to examine newly synthesized β-actin specifically, we find a significant increase in growth cones of neurons cultured on laminin 111 in the presence of BDNF. However, soluble laminin 111 alone results in a significant reduction in nascent β–actin protein synthesis. These results indicate that laminin isoforms can act in multiple ways, including synergistically with guidance cues and independently, to modulate local mRNA translation, thereby differentially influencing axon growth and guidance during development.

**Summary Statement:** Local translation in axons is critical for axon guidance. Laminin, a key component of the extracellular matrix, is necessary to induce local translation and thus mediate axon growth and guidance.

## Introduction

Accurate neural wiring is important for the formation of a healthy, functional brain. Changes in neural wiring or the failure to connect with synaptic targets can give rise to various neurological disorders (Van Battum et al., 2015). During development, this neural network is formed through axon guidance, wherein neuronal processes are directed to and establish connections with their synaptic targets (Bellon and Mann, 2018). Axon guidance is mediated by growth cones, which are highly dynamic and motile motor and sensory structures located at the tips of pathfinding axons. Growth cones respond to extracellular cues in their environment, which can be attractive or repulsive, as well as diffusible or contact-mediated (Gomez et al., 1996, Lowery and Vactor, 2009, Bixby and Harris, 1991, McFarlane and Holt, 1997). These cues are sensed by the receptors present on growth cone filopodia and lamellipodia, initiating signaling mechanisms that reorganize the cytoskeleton and allow the growth cone to advance towards, stall, or turn away from the cue (Myers et al., 2011).

Contact-mediated cues, such as extracellular matrix (ECM) proteins, are critical in axon guidance. Laminin is a major component of the ECM and is widely expressed in both the peripheral and central nervous systems (Barros et al., 2011, McKerracher et al., 1996, Myers et al., 2011). Numerous studies have reported that laminin regulates axon guidance (Barros et al., 2011, Kuhn et al., 1995, McKerracher et al., 1996, Bonner and O’Connor, 2001, Paulus and Halloran, 2006). Moreover, netrin-1 is an attractive guidance cue for retinal neurons, but when a high concentration of laminin substrate is also present, netrin-1 becomes repellent to these neurons (Hopker et al., 1999). Similarly, retinal ganglion cells collapse in the presence of EphB and laminin, but when L1 is also present, then growth cone pausing occurs (Suh et al., 2004). Thus, laminin acts in concert with other guidance molecules to differentially remodel the cytoskeleton, but we currently have limited knowledge about how this signaling from multiple cues is integrated.

One mechanism critical for axon guidance is local translation, the process by which mRNAs are localized to specific subcellular locations and rapidly translated on-site in response to extracellular guidance cues (Van Battum et al., 2015, Hengst and Jaffrey, 2007, Lin and Holt, 2007). Thousands of mRNAs are localized to developing axons (Taylor et al., 2009, Gumy et al., 2011, Zivraj et al., 2010, Lin et al., 2021). Among these are mRNAs encoding cytoskeletal proteins, such as β-actin, that are a fundamental part of the growth cone structure and dynamically altered to mediate axon guidance (Bassell et al., 1998, Yao et al., 2006). It has been shown that brain-derived neurotrophic factor (BDNF) and netrin-1-induced local translation of *β-actin* mRNA in growth cones is necessary for attractive turning of the growth cone and thus for axon guidance (Yao et al., 2006, Leung et al., 2006a, Sasaki et al., 2010, Kershner et al., 2019, Welshhans and Bassell, 2011). Additionally, mRNAs of axon growth-promoting proteins, such as GAP43, are locally translated in response to attractive guidance cues like nerve growth factor (Donnelly et al., 2013, Smith et al., 2004). Similarly, repulsive cues, such as Sema3A and Slit2, can locally translate RhoA and cofilin, leading to growth cone collapse or depolymerization of actin filaments, respectively (Dent et al., 2011, Piper et al., 2006, Wu et al., 2005). This shows that the nature of guidance cues can dictate which mRNA subsets are translated.

Although guidance cues are well known to regulate local translation, almost nothing is known about the role of the ECM in this process. In non-neuronal cells, the substrate can influence the pool of mRNAs that are translated (Benecke et al., 1978, Farmer et al., 1978). Furthermore, one study in embryonic mouse motoneurons has shown that local translation of *β-actin* mRNA in growth cones is induced by laminin 111 (Rathod et al., 2012), a widely expressed isoform of laminin, both spatially and temporally (Miner et al., 1997, Lentz et al., 1997, Libby et al., 2000, Liesi et al., 2001, Sasaki et al., 2002, Nirwane and Yao, 2019). In contrast, embryonic motoneurons cultured on laminin 211/221 showed a significant reduction in *β-actin* mRNA local translation (Rathod et al., 2012). This is critical as it is the first and only study to examine the role of laminin in local translation.

Other well-established roles of laminin include adhesion formation and neurite outgrowth (Luckenbill-Edds, 1997, Plantman et al., 2008, Woo and Gomez, 2006, Nichol et al., 2016, Murtomaki et al., 1992), which is mediated through the classical laminin receptors, integrins (Plantman et al., 2008, Lathia et al., 2007, Lei et al., 2012). Adhesions in growth cones are termed point contacts. Point contacts are dynamic multiprotein adhesion complexes that are composed of integrins, paxillin, talin, vinculin, and RACK1 (Renaudin et al., 1999, Chastney et al., 2021, Kershner and Welshhans, 2017). Point contacts link the ECM to the intracellular actin cytoskeleton via transmembrane integrin receptors and restrain the retrograde flow of actin, allowing increased polymerization at the barbed end, and thus facilitating membrane protrusion (Nichol et al., 2016, Gallo, 2011, Gomez and Letourneau, 2014). Moreover, they are critical for axon growth and guidance, as disruption of point contact formation and turnover impairs axon outgrowth and growth cone response to guidance cues (Woo and Gomez, 2006, Robles and Gomez, 2006, Myers and Gomez, 2011, Becret et al., 2025). Nichol et al previously showed that dynamic point contacts are increased, while retrograde flow is reduced when neurons are cultured on laminin (Nichol et al., 2016).

This is interesting as our lab has previously shown that RACK1, *β-actin* mRNA, and ribosomal subunits are localized at point contacts and that local translation of *β-actin* occurs at these adhesions (Kershner et al., 2019, Kershner and Welshhans, 2017). Thus, point contacts are an optimal site for integrating guidance-cue signaling via local translation, thereby directly mediating axon guidance.

Here, we investigated whether laminin and BDNF have distinct or synergistic signaling roles in regulating local translation within growth cones. Our findings show that laminin 111, when presented as a substrate or acutely in the presence of BDNF, synergistically increases local translation of mRNAs in mouse embryonic day 17 cortical neurons. However, laminin isoforms 211 and 221 inhibit local translation at growth cones in the presence of BDNF. We also show that BDNF and substrate laminin 111 increase the local translation of *β-actin* mRNA, whereas laminin alone increases the number of growth cones actively translating this protein. Finally, we also report that acute addition of soluble laminin decreases local translation of *β-actin* mRNA. These results provide new insights into how ECM proteins coordinate signaling with diffusible guidance cues to regulate local translation, subsequently affecting axon guidance and neural circuit formation.

## Results

### Laminin increases BDNF-induced local translation in growth cones

Neurotrophic factors, such as BDNF and Netrin-1, induce local translation in axons (Takei et al., 2004, Ruiz et al., 2014, Jain and Welshhans, 2016, Welshhans and Bassell, 2011, Leung et al., 2006a). Most studies examining local translation have cultured neurons on a laminin substrate; however, only one study has examined whether laminin affects local translation (Rathod et al., 2012). Because laminin regulates axon guidance (Hopker et al., 1999, Bonner and O’Connor, 2001, McKerracher et al., 1996), we investigated whether laminin affects local translation in growth cones of developing mouse cortical neurons. Here, we cultured E17 mouse cortical neurons on one of two substrates: (1) poly-L-lysine (PLL) alone, or (2) PLL + Laminin 111. In both conditions, we performed a starve and stimulate paradigm with BDNF at 2 DIV (**Fig. 1A**). Neurons were stimulated with BDNF for 10 minutes, followed by a 10-minute incubation with both BDNF and puromycin. Puromycin is an analog of tRNA that incorporates into actively elongating polypeptide chains; following fixation, puromycin can be detected using immunofluorescence, thus labeling nascent peptide chains (tom Dieck et al., 2015). To confirm the known growth-promoting effects of laminin on cultured neurons, we first quantified axon length. Neurons cultured on PLL+ laminin and stimulated with BDNF showed significantly increased axon length compared to those cultured on PLL and stimulated with BDNF (**Fig. 1B and 1C**). However, quantification of growth cone area showed no differences between the two groups (**Fig. 1D**).

**Figure 1:**
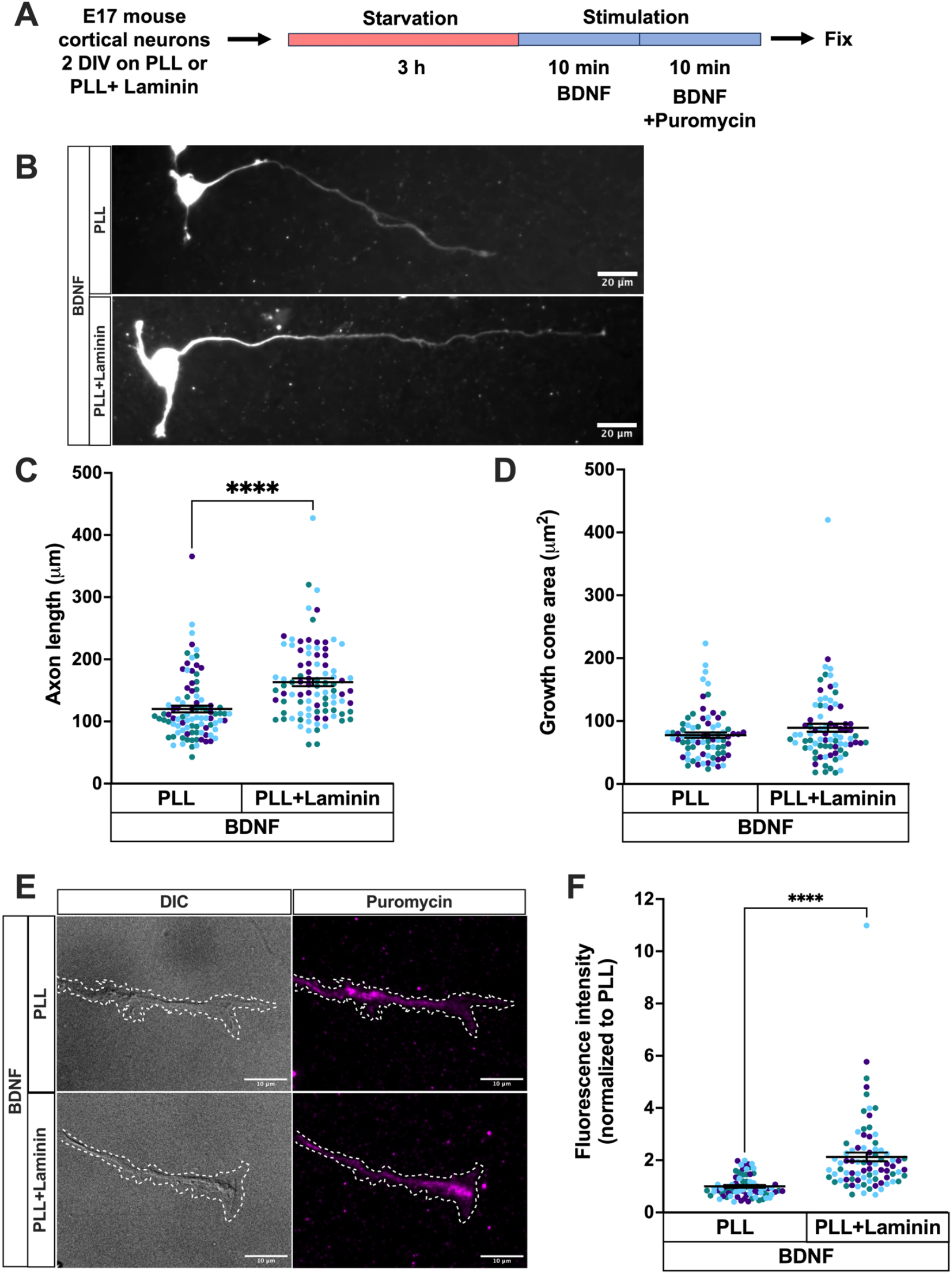
Laminin increases BDNF-induced local translation. **(A)** Primary E17 mouse cortical neurons were cultured for 2 days *in vitro* on either poly-l-lysine (PLL) alone or PLL and mouse Laminin 111. Following 2 DIV in culture, neurons were starved of growth factors for 3 h. Next, stimulation media containing growth factors and BDNF were added for 10 min, then puromycin was added to the stimulation media for 10 min, followed by immediate fixation. **(B)** Representative images of E17 cortical neurons immunostained for puromycin. Scale bar, 20μm. **(C)** Axon length was quantified. Two-tailed Mann-Whitney test, *\*\*\*\*p*<0.0001. PLL + BDNF, n = 92, and PLL + Laminin + BDNF, n = 92, across 3 separate experiments, plotted as three separate colors. **(D)** Growth cone area was quantified. Two-tailed Mann-Whitney test, *p*=0.288. PLL + BDNF, n = 79, and PLL + Laminin + BDNF, n = 80, across 3 separate experiments, plotted as three separate colors. **(E)** Representative images (DIC and epifluorescence) of growth cones immunostained for puromycin (magenta). The dotted line outlines both growth cones and axons for ease of visualization; however, axons were not included in the analysis. Scale bars, 10 μm. **(F)** Puromycin intensity was quantified in neuronal growth cones. Two-tailed Mann-Whitney test, *\*\*\*\* p*<0.0001, PLL + BDNF, n = 79, and PLL + Laminin + BDNF, n = 80, across 3 separate experiments, plotted as three separate colors.

To examine local translation, we measured the fluorescence intensity of puromycin in growth cones of these cortical neurons. Growth cones of neurons cultured on PLL + Laminin 111 and stimulated with BDNF showed a significant increase in local translation compared to those cultured on PLL alone and stimulated with BDNF (**Fig. 1E and 1F**). Thus, these results suggest that the ECM protein laminin stimulates BDNF-induced local translation in growth cones.

### Laminin isoforms differentially impact local translation

Laminin isoform expression in the CNS and PNS is spatiotemporally regulated across development and into adulthood (Lathia et al., 2007, Liesi and Silver, 1988, Nirwane and Yao, 2019, Rogers and Nishimune, 2017, Nirwane and Yao, 2022, Copp et al., 2011). Laminins 111 and 511 are the first isoforms expressed during development and are continually expressed until adulthood (Nirwane and Yao, 2019, Copp et al., 2011, Miner et al., 2004). Other laminin isoforms, such as 211, are reported to be secreted by glial cells and present in skeletal muscles (Liesi and Silver, 1988, Miner et al., 2004), while laminin 221 is expressed at the neuromuscular junction (Rogers and Nishimune, 2017). A pioneering study examining the role of laminins in local translation showed that laminin 111, in the presence of BDNF, increased *β*-actin local synthesis in mouse motoneurons compared with a mix of laminin 211/221 (Rathod et al., 2012). Given this finding, we examined how a variety of laminin isoforms affect local translation in cortical neurons, both in the presence and absence of BDNF. We cultured E17 cortical neurons on PLL alone or PLL plus human recombinant laminin 111, 511, 211, or 221 for 2 DIV, then performed a starve and stimulate paradigm with BDNF and a puromycin assay (**Figure 2A**). The fluorescence intensity of puromycin was quantified in growth cones. Local translation in growth cones cultured on human recombinant laminin 111 or 511 was not significantly different as compared to those cultured on PLL. However, local translation was significantly decreased in growth cones of neurons cultured on laminins 211 or 221 compared with those cultured on PLL (**Fig. 2B and C**).

**Figure 2:**
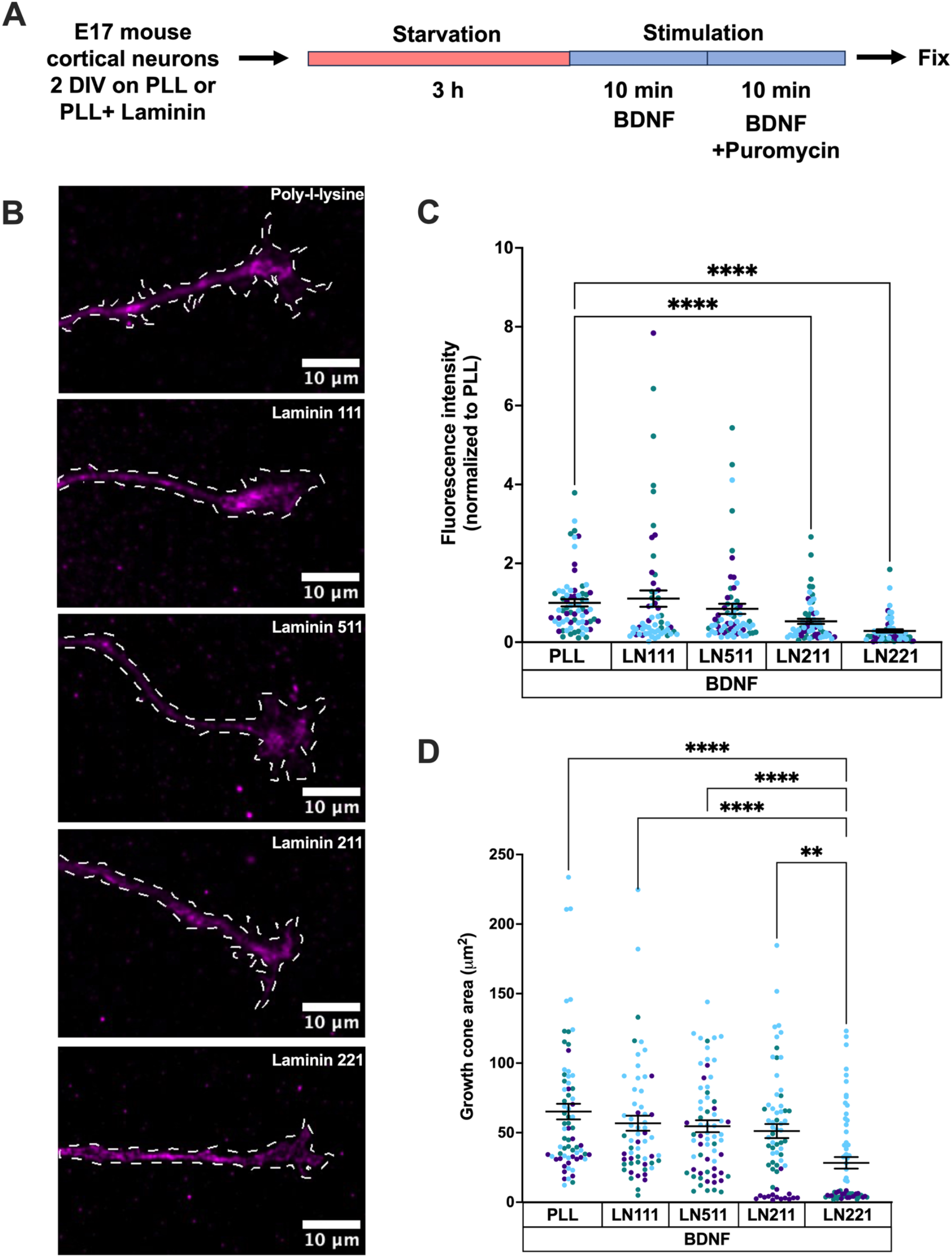
Laminins 211 and 221 significantly decrease BDNF-induced local translation. **(A)** Primary E17 mouse cortical neurons were cultured on PLL, PLL + Laminin 111, PLL + Laminin 511, PLL + Laminin 211, or PLL + Laminin 221 for 2 DIV. Neurons were stimulated with BDNF via bath application for 10 min. Next, puromycin was added to the stimulation media for 10 min, and cells were fixed immediately. **(B)** Representative images of growth cones immunostained for puromycin (magenta). The dotted line outlines both growth cones and axons for ease of visualization; however, axons were not included in the analysis. Scale bars, 10 μm. Puromycin intensity **(C)** and growth cone area **(D)** were quantified in neuronal growth cones. ANOVA, Dunn’s Post-hoc test, \*\**p* < 0.01 and \**\*\*\*p* < 0.0001. PLL + BDNF, n = 69; Laminin 111 + BDNF, n = 58; Laminin 511 + BDNF, n = 65; Laminin 211 + BDNF, n = 65; Laminin 221 + BDNF, n = 64, across 3 separate experiments, plotted as three separate colors.

As local translation was significantly inhibited by laminin 211 or 221, we quantified growth cone area. Neurons cultured on PLL + Laminin 221 have significantly smaller growth cone area compared to all the other groups, but growth cone size was not affected by laminin 211 (**Fig. 2D**). These data indicate that laminin isoforms can differentially regulate local translation and growth cone morphology.

### Laminin and BDNF co-regulate local translation of *β-actin* mRNA in growth cones

We next sought to identify specific mRNAs that would be locally translated in response to laminin and BDNF. In both non-neuronal and neuronal cells, *β-actin* mRNA has been studied extensively and is known to be localized and translated in the leading edge of fibroblasts and in neuronal growth cones in response to growth and guidance cues (Welshhans and Bassell, 2011, Bassell et al., 1998, Yao et al., 2006, Condeelis and Singer, 2005, Hill and Gunning, 1993, Lawrence and Singer, 1986, Leung et al., 2006b). Thus, we examined the role of mouse laminin 111 in the local translation of *β-actin* mRNA. E17 mouse cortical neurons were cultured on PLL alone or PLL + laminin 111 for 2 DIV, followed by a starve and stimulate paradigm with BDNF (or vehicle) and a puromycin assay (**Fig. 3A**). After nascent polypeptides were puromycylated, a puromycin-proximity ligation assay (Puro-PLA; **Fig. 3B**) was used to fluorescently label locally synthesized β-actin protein, visible as individual puncta within growth cones (**Fig. 3C**). The fluorescent intensity of individual puncta within growth cones was quantified using a particle analysis tool in Fiji. Growth cones without any puncta were excluded from the puncta intensity analysis. PLL + BDNF or laminin 111 + Vehicle (for BDNF) were not sufficient to significantly increase the amount of β-actin locally synthesized in the growth cones compared to the control group (Vehicle + PLL). However, when neurons were cultured on laminin 111 and stimulated with BDNF, the local synthesis of β-actin in growth cones was significantly increased (**Fig. 3C and 3D**). These data suggest that either BDNF alone or laminin 111 alone is not sufficient to stimulate the local translation of β-actin; rather, both BDNF and laminin 111 are needed for this local translation.

**Figure 3:**
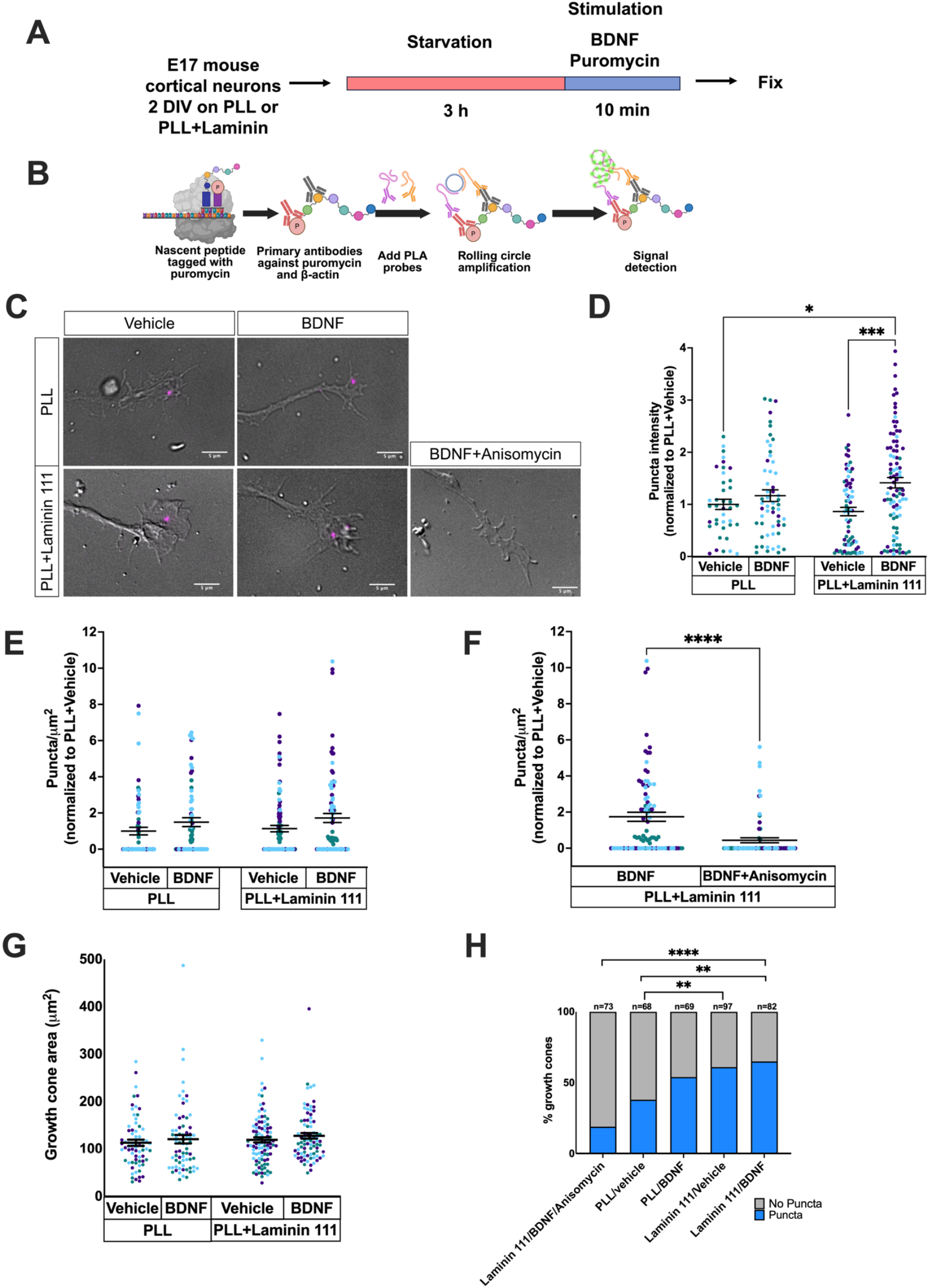
LN111 and BDNF co-regulate local translation of *β-actin* mRNA. **(A)** E17 mouse cortical neurons were cultured on PLL or PLL + Laminin 111 for 2 DIV, then starved for 3 h. Neurons were then stimulated with either vehicle for BDNF or BDNF along with puromycin via bath application for 10 min. Cells were fixed immediately. **(B)** Puromycin-proximity ligation assay (Puro-PLA) was performed after fixation to label nascent β-actin protein. Primary antibodies to puromycin and β-actin are added, following which secondary antibodies with PLA plus and minus probes bind to the primary antibodies. When within 40 nm of each other, the probes initiate rolling circle amplification upon addition of ligase and fluorescently labeled dNTPs, yielding a fluorescent detection signal for the protein of interest, here β-actin. **(C)** Representative images of β-actin puro-PLA (magenta) signal overlaid on the corresponding DIC image. Scale bars 5 μm. **(D)** Quantification of β-actin puro-PLA puncta intensity in growth cones. Growth cones showing no puncta were excluded from the puncta intensity analysis. Two-way ANOVA, Tukey’s multiple comparisons, *p<0.05 and ***p<0.0001. PLL + vehicle, n = 39; PLL + BDNF, n = 56; Laminin 111 + vehicle, n = 72; Laminin 111 + BDNF, n = 92, where n= total number of puncta measured across 3 separate experiments. **(E, F)** Quantification of the density of locally synthesized β-actin puncta was calculated as the number of β-actin puncta per growth cone divided by the area of the growth cone. (E) Two-way ANOVA, Tukey’s multiple comparisons; PLL + vehicle, n = 68; PLL + BDNF, n = 69; Laminin 111 + vehicle, n = 97; Laminin 111 + BDNF, n = 82, where n= number of growth cones measured across 3 separate experiments. (F) Two-tailed Mann-Whitney test, ***p=0.003; Laminin111 + BDNF, n = 82; Laminin111 + BDNF + Anisomycin, n = 73, where n= total number of growth cones measured across 3 separate experiments. **(G)** Quantification of growth cone area. Two-way ANOVA, Tukey’s multiple comparisons; PLL + vehicle, n = 68; PLL + BDNF, n = 69; Laminin 111 + vehicle, n = 97; Laminin 111 + BDNF, n = 82, where n= number of growth cones measured across 3 separate experiments. **(H)** Percentage of growth cones showing puncta versus growth cones showing no puncta. Chi-square test (*p* < 0.0001). Bonferroni’s correction was applied to adjust for multiple comparisons, and the p-value for post-hoc tests was set at 0.0083. PLL + Vehicle vs PLL + BDNF, p = 0.078; PLL + BDNF vs PLL + BDNF + Anisomycin, p< 0.0001; PLL + Vehicle vs Laminin + Vehicle, p = 0.0043; PLL + BDNF vs Laminin + BDNF, p = 0.169; Laminin + Vehicle vs Laminin + BDNF, p = 0.599; Laminin + BDNF vs Laminin + BDNF + Anisomycin, p < 0.0001. PLL + vehicle, n = 68; PLL + BDNF, n = 69; Laminin 111 + Vehicle, n = 97; Laminin 111 + BDNF, n = 82; PLL + BDNF +Anisomycin, n = 63; Laminin 111 + BDNF +Anisomycin, n = 73, n= number of growth cones measured across 3 separate experiments.

β-actin puncta density, quantified as the number of β-actin puro-PLA puncta per growth cone divided by the area of the growth cone, was not significantly different between any experimental condition (**Fig. 3E**). However, treatment of neurons cultured on laminin 111, pretreated with the protein synthesis inhibitor anisomycin, and stimulated with BDNF showed a significant decrease in β-actin puncta density (**Fig. 3F**), confirming effective inhibition of protein synthesis and the validity of our Puro-PLA. No significant differences in growth cone area were observed between conditions (**Fig. 3G**). Additionally, we quantified the percentage of growth cones with or without any β-actin puro-PLA puncta in each group, as a measure of the number of growth cones engaged in local translation. Laminin 111 + Vehicle and Laminin 111 + BDNF resulted in a significantly higher number of growth cones with β-actin puncta, as compared to the PLL + Vehicle group (**Fig. 3H**). There was no significant difference between the Laminin 111 + Vehicle and the Laminin 111 + BDNF groups (**Fig. 3H**); these data suggest that laminin may actively engage more growth cones in *β-actin* mRNA local translation. Taken together, these data indicate that both laminin 111 and BDNF are needed to increase *β-actin* mRNA local translation within growth cones, but laminin 111 alone regulates which growth cones are actively engaged in local translation.

### Acutely encountering laminin 111 and BDNF synergistically increases paxillin-rich point contact density, axon length, and local translation in growth cones

In the embryonic brain, growing axons encounter laminin in subregions of their migration tract, the expression of which is tightly regulated by the developmental timelines (Nirwane and Yao, 2019, Libby et al., 2000, Miner et al., 1997, Lentz et al., 1997, Indyk et al., 2003). This suggests that axonal growth cones are not continually exposed to laminin. Moreover, laminin expression levels and deposition patterns vary spatiotemporally (Lathia et al., 2007, Miner et al., 2004, Miner et al., 1997, Lentz et al., 1997, Hamill et al., 2009). In Figure 1-3, neurons were cultured on a low concentration of laminin (10 μg/mL) for the entirety of two days. Here, we modified our experimental design to more closely mirror what may occur in the developing brain. E17 mouse cortical neurons were cultured on a PLL-only substrate for 2DIV. Next, we performed a modified starve and stimulate paradigm. This consisted of starvation for 3h, followed by acute stimulation with one of 4 conditions for 25 minutes: (1) Vehicles (Vehicle for BDNF + Vehicle for Laminin 111), (2) BDNF alone (BDNF + Vehicle for Laminin 111), (3) 25 μg/mL Laminin 111 alone (Vehicle for BDNF + Laminin 111), or (4) BDNF and 25 μg/mL Laminin 111 (BDNF + Laminin 111). Puromycin was added during the last 10 minutes of the stimulation period, and then neurons were fixed (**Fig. 4A**). The timeline was adjusted in this experiment to account for the time required for the laminin to adhere to PLL (Nichol et al., 2016). Given that substrate laminin promotes axon outgrowth and point contact formation (Robles and Gomez, 2006, Myers and Gomez, 2011, Cohen et al., 1987, Murtomaki et al., 1992, Plantman et al., 2008), we measured axon length and paxillin-rich point contact density to examine the effect of soluble laminin. PLL (Vehicle for BDNF + Vehicle for Laminin 111), BDNF alone (BDNF + Vehicle for Laminin 111), or Laminin 111 alone (Vehicle for BDNF + Laminin 111) were not enough to cause a significant increase in point contact density (paxillin density), local translation (puromycin intensity), or axon length (**Fig. 4B-E, and G-H**). However, acute stimulation with BDNF + laminin 111 resulted in a significant increase in point contact density, local translation (puromycin intensity), and axon length, as compared to all other groups. Growth cone area was significantly larger in neurons stimulated with laminin 111 alone (Vehicle for BDNF + Laminin 111) than those treated with both laminin 111 + BDNF (**Fig. 4F**). Taken together, our data suggest that axons encountering laminin 111, either as an evenly applied substrate or as an acute stimulant, increases local translation of mRNAs in the presence of BDNF.

**Figure 4:**
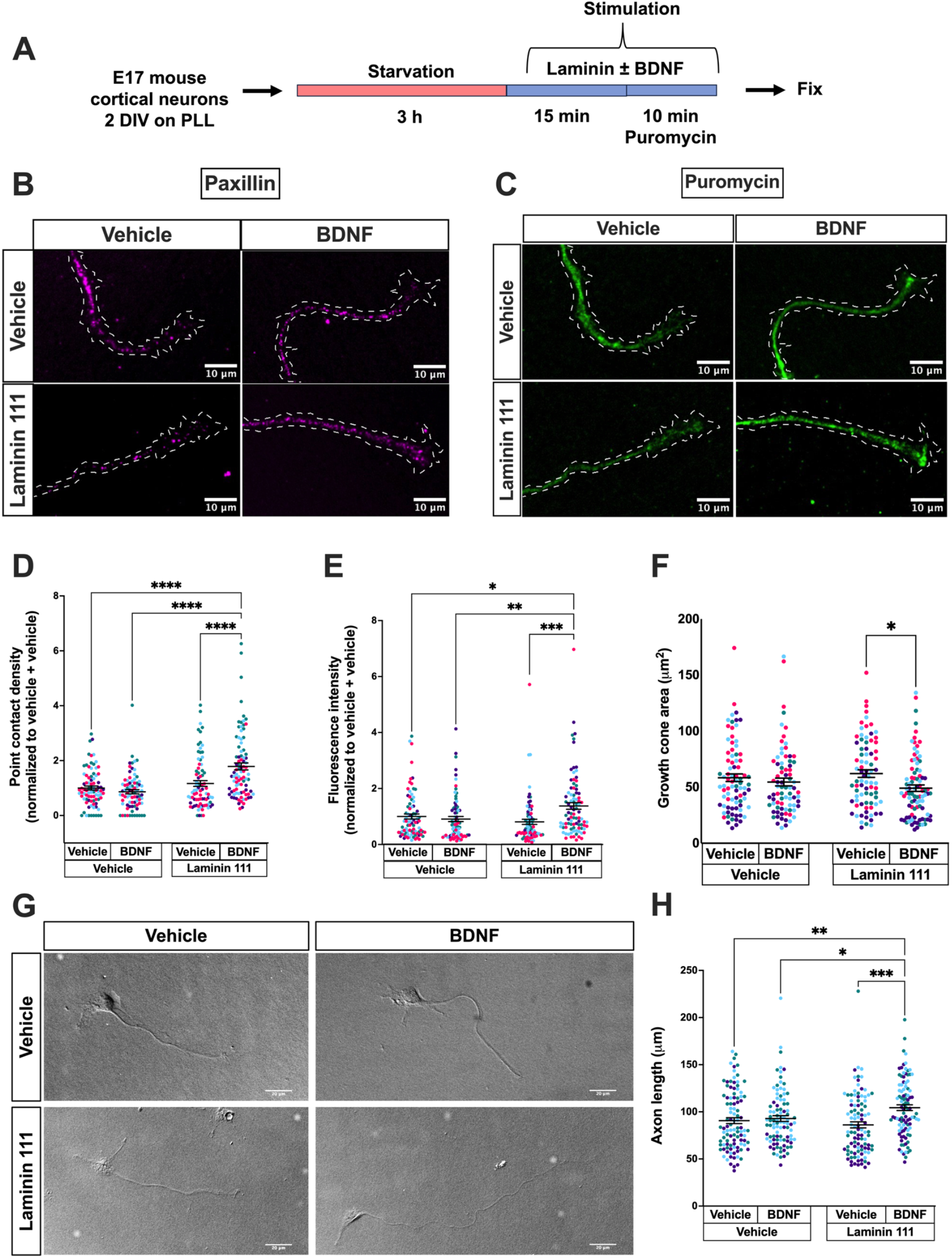
Acutely encountering Laminin 111 and BDNF synergistically increases point contact density, axon growth, and local translation in growth cones. **(A)** Primary E17 mouse cortical neurons were cultured on PLL for 2 DIV. Neurons were then starved of growth factors for 3 h. Next, neurons were stimulated with either: (1) Vehicles (Vehicle for BDNF/Vehicle for Laminin 111), (2) BDNF alone (BDNF/Vehicle for Laminin 111), (3) Laminin 111 alone (Vehicle for BDNF/Laminin 111), or (4) BDNF and Laminin 111 (BDNF/Laminin 111) in a bath application for 15 min. Next, puromycin was added to the stimulation media for 10 min, and cells were fixed immediately. **(B, C)** Representative images of growth cones immunostained for paxillin, a point contact marker (magenta; B), and puromycin (green; C). Dotted line shows growth cones and the axon shaft for ease of visualization, but axons were not included in the analysis. Scale bars, 10 μm. **(D-F)** Quantification of point contact density, using paxillin as a point contact marker (D), puromycin intensity (E), and growth cone area (F). Two-way ANOVA, Tukey’s multiple comparisons, \**p* < 0.05, \*\**p* < 0.01, \*\*\**p* < 0.001, \*\*\*\**p* < 0.0001; vehicle + vehicle, n = 83; vehicle + BDNF, n = 78; Laminin 111 + vehicle, n = 80; Laminin 111 + BDNF, n = 90, across 4 separate experiments. **(G)** Representative DIC images used to quantify axon length. Scale bars, 20 μm. **(H)** Quantification of axon length. Two-way ANOVA, Tukey’s multiple comparisons, \**p* < 0.05, \*\**p* < 0.01, \*\*\*\**p* < 0.0001; vehicle + vehicle, n = 101; vehicle + BDNF, n = 94; Laminin 111 + vehicle, n = 99; Laminin 111 + BDNF, n = 99, across 3 separate experiments

### Acutely encountering laminin 111 decreases local translation of *β-actin* mRNA

We now repeated the acute encounter paradigm and treatment groups used in Figure 4 to determine if the local translation of *β-actin* mRNA specifically was synergistically regulated by soluble laminin 111 and BDNF. We performed a Puro-PLA for β-actin after labeling all nascent peptides with puromycin (**Fig. 5A**), and measured puncta intensity, puncta density, growth cone area, and percentage of growth cones with β-actin puncta, as described in Figure 3. Interestingly, the intensity of β-actin puncta was significantly decreased when stimulated with soluble laminin 111 alone (Vehicle for BDNF + Laminin 111), as compared to vehicle (Vehicle for BDNF + Vehicle for Laminin 111) or BDNF alone (BDNF + Vehicle for Laminin 111) groups (**Fig. 5B and C**). Moreover, BDNF alone (BDNF + Vehicle for Laminin 111) had significantly higher *β-actin* mRNA local translation compared to neurons exposed to BDNF and laminin together (BDNF + Laminin 111) (**Fig. 5B and C**). For β-actin puncta density, there was a significant increase in neurons treated with BDNF and laminin together (BDNF + Laminin 111), as compared to laminin alone (Vehicle for BDNF + Laminin 111) (**Fig. 5D**). The negative control, anisomycin, effectively inhibited translation, showing specificity of the puro-PLA signal (**Fig. 5E**). There were no significant differences in growth cone area across the various groups (**Fig. 5F**). Finally, unlike the effect of substrate laminin increasing the percentage of growth cone actively involved in local translation (**Fig. 3H**), acutely applying laminin resulted in no significant difference between the groups, except that the anisomycin group showing a significantly lower percentage of growth cones translating *β-actin* mRNA, as expected (**Fig. 5G**). These findings reveal that acute encounters with soluble laminin 111 may activate distinct signaling pathways, as compared to substrate laminin, to affect *β-actin* mRNA local translation.

**Figure 5:**
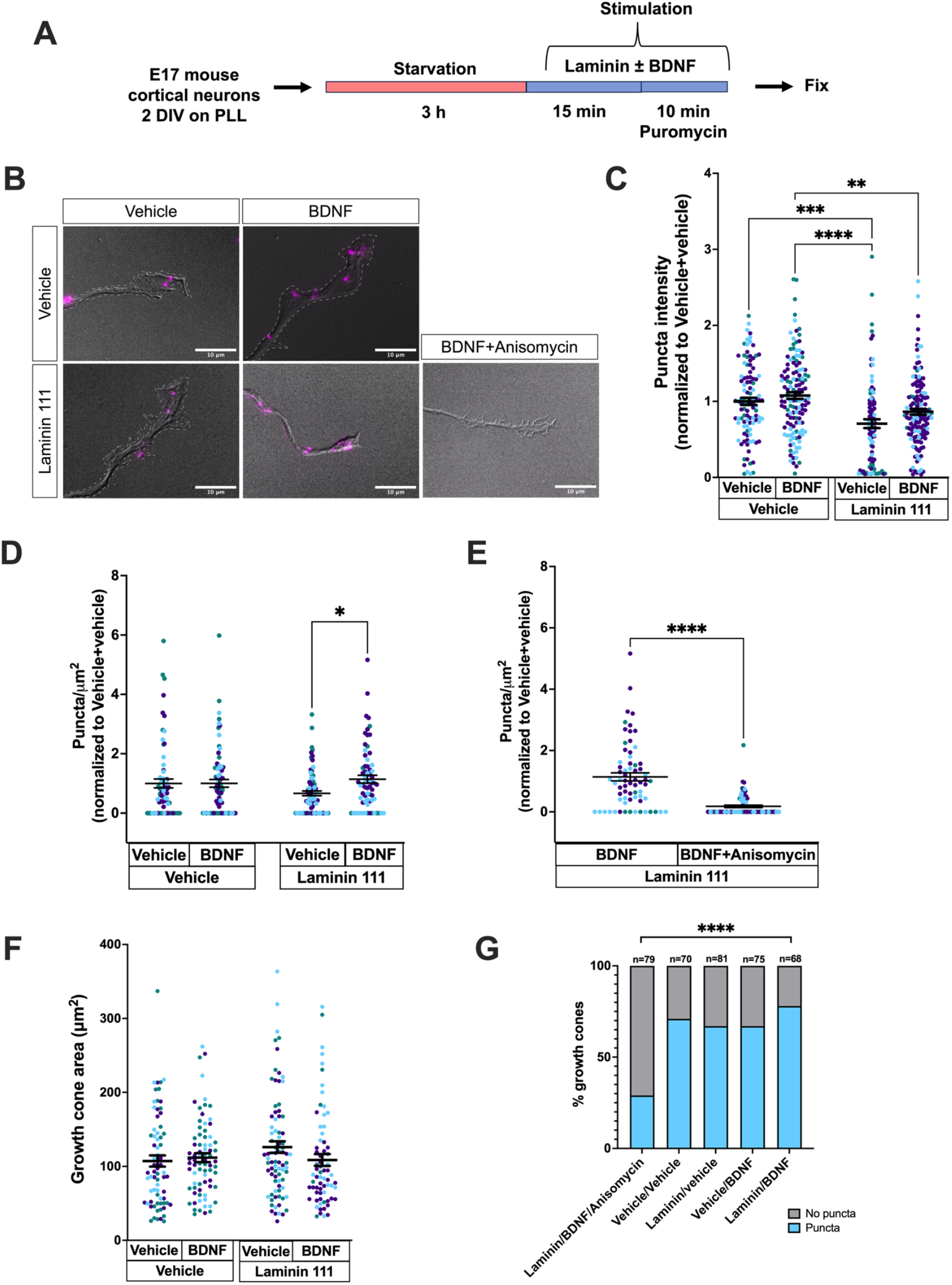
Acutely encountering laminin 111 decreases β-actin local translation. **(A)** Primary E17 mouse cortical neurons were cultured on PLL for 2 DIV. Neurons were starved of growth factors for 3 h and then stimulated with either: (1) Vehicles (Vehicle for BDNF/Vehicle for Laminin 111), (2) BDNF alone (BDNF/Vehicle for Laminin 111), (3) Laminin 111 alone (Vehicle for BDNF/Laminin 111), or (4) BDNF and Laminin 111 (BDNF/Laminin 111) in a bath application for 15 min. Next, puromycin was added to the stimulation media for 10 min, and then cells were fixed immediately. A puromycylation proximity ligation assay (Puro-PLA) was performed after fixation to label nascent β-actin protein. **(B)** Representative images of β-actin puro-PLA (magenta) signal overlaid on the corresponding DIC image. Scale bars, 10 μm. **(C)** Quantification of β-actin puro-PLA puncta intensity in growth cones. Growth cones showing no puncta were excluded from the puncta intensity analysis. Scale bars 10 μm. Two-way ANOVA, Tukey’s multiple comparisons, \*\**p* < 0.01, \*\*\**p* < 0.001, \*\*\*\**p* < 0.0001; vehicle + vehicle, n = 105; vehicle + BDNF, n = 134; Laminin 111 + vehicle, n = 96; Laminin 111 + BDNF, n =142, where n is the total number of puncta measured across 3 separate experiments. **(D, E)** Quantification of the density of locally synthesized β-actin puncta was calculated as the number of β-actin puncta per growth cone divided by the area of the growth cone. (D) Two-way ANOVA, Tukey’s multiple comparisons, \**p* < 0.05; vehicle + vehicle, n = 70; vehicle + BDNF, n = 75; Laminin 111 + vehicle, n = 81; Laminin 111 + BDNF, n =68, where n is the total number of growth cones measured across 3 separate experiments. (E) Two-tailed Mann-Whitney test, \*\*\*\**p* < 0.0001; Laminin 111 + BDNF, n = 68; Laminin 111 + BDNF + Anisomycin, n = 79, where n is the total number of growth cones measured across 3 separate experiments. **(F)** Quantification of growth cone area. Two-way ANOVA, Tukey’s multiple comparisons; Vehicle + vehicle, n = 70; Vehicle + BDNF, n = 75; Laminin 111 + vehicle, n = 81; Laminin 111 + BDNF, n =68, where n is the total number of growth cones measured across 3 separate experiments **(G)** Percentage of growth cones showing puncta versus growth cones showing no puncta. Chi-square test (*p* < 0.0001). Bonferroni’s correction was applied to adjust for multiple comparisons, and the p-value for post-hoc tests was set at 0.01. vehicle + vehicle vs vehicle + BDNF, p = 0.5408; vehicle + vehicle vs Laminin + vehicle, p = 0.5408; vehicle + vehicle vs Laminin +BDNF, p=0.2561; vehicle + BDNF vs Laminin + BDNF, p = 0.0815; Laminin + vehicle vs Laminin + BDNF, p = 0.0815; Laminin + BDNF vs Laminin + BDNF + Anisomycin, p < 0.0001. Vehicle + vehicle, n = 70; vehicle + BDNF, n = 75; Laminin 111 + vehicle, n = 81; Laminin 111 + BDNF, n = 68; Laminin 111 + BDNF +Anisomycin, n = 79, where n is the total number of growth cones measured across 3 separate experiments.

## DISCUSSION

Laminin is necessary for multiple aspects of neural development, including adhesion formation, neurite outgrowth, and axon guidance (Gomez et al., 1996, Hantaz-Ambroise et al., 1987, Cohen and Johnson, 1991, Clark et al., 1993, Hopker et al., 1999, Myers et al., 2011, Woo and Gomez, 2006, Nichol et al., 2016). Although local translation is a critical mechanism for axon growth, maintenance, and guidance (Smith et al., 2004, Donnelly et al., 2013, Welshhans and Bassell, 2011, Cox et al., 2008, Martin, 2004, Spillane et al., 2013), to the best of our knowledge, only one study has examined the role of laminin in local translation (Rathod et al., 2012). Using a fluorescent *β-actin* translation reporter and examining the axons of mouse motor neurons, this study demonstrated that *β-actin* mRNA is locally translated in neurons cultured on laminin 111, but is significantly reduced in neurons cultured on a laminin substrate containing a mix of both laminin 211 and 221. In both of these experiments, BDNF was included in the culture media. Interestingly, this process is reversed in SMN (survival of motoneuron)-deficient motoneurons. This study provided an important foundation for the current work, but it did not examine the discrete contributions of laminin and BDNF to local translation. The current study fills that important gap in the literature. Furthermore, our study examines the sufficiency of laminin and BDNF for both generalized local translation (i.e., puromycin assay) and the local translation of specific mRNAs (i.e., *β-actin*). Finally, the current study more closely replicates how axons may encounter laminin *in vivo* in the developing brain (Zhou, 1990, Luckenbill-Edds, 1997) and examines how the concentration of laminin affects axonal local translation.

Our study is the first to show that laminin is necessary for BDNF-induced local protein synthesis in growth cones of mouse embryonic cortical neurons; however, the effect of laminin on local translation depends on its concentration and how neurons encounter it. We show that low concentrations (10 μg/mL) of substrate laminin, and high concentrations (25 μg/mL) of soluble laminin increased BDNF-induced local translation in growth cones (Fig.1E,1F, 4C, and 4E). This suggests that growth cones that continually encounter laminin over two days and are then acutely exposed to BDNF, or that are acutely exposed to both laminin and BDNF simultaneously via bath application, integrate these cues to produce a synergistic effect. This finding that laminin stimulates local translation is particularly important because there are many papers demonstrating that neurotrophic factors, such as BDNF, netrin-1, and NT-3, induce local translation (Takei et al., 2004, Ruiz et al., 2014, Jain and Welshhans, 2016, Welshhans and Bassell, 2011, Spillane et al., 2012, Leung et al., 2006a, Koppers et al., 2024). These studies used laminin-coated substrates in their experiments, as it is routinely used for culturing many neuronal types, but none examined whether laminin is necessary for this molecular mechanism. The current study is the first to demonstrate that neither laminin alone nor BDNF alone is sufficient to increase local translation; synergistic signaling is required to induce this important molecular mechanism.

The role of laminin in regulating local translation can be further elucidated by examining its isoforms. In our study, we showed that human recombinant isoforms (LN111, LN511, LN211, and LN221) used at a low concentration (10 μg/mL) as substrate have differential effects on local translation (Fig. 2). While we did not see a significant increase with human recombinant laminins 111 and 511, laminins 211 and 221 inhibited BDNF-induced local translation (Fig. 2B and C). The amino acid sequence of mouse and human laminins differs by about 30% (Barraza-Flores et al., 2020), and thus, it is likely that mouse neurons interact with mouse origin laminin 111 more effectively than the human recombinant laminins 111 and 511 used in our study, which may not initiate signaling. Although laminins 211 and 221 were also of human origin, these two laminins are typically enriched in glial cells, skeletal muscle, or at neuromuscular junctions (Holmberg and Durbeej, 2013, Liesi and Risteli, 1989, Hall, 1995, Sasaki et al., 2002, Nirwane and Yao, 2019, Liesi et al., 2001) and may have higher similarity to mouse (Durkin et al., 1996). Thus, the binding of these isoforms to mouse neuron membrane receptors is likely effective. Unfortunately, other than laminin 111, full-length recombinant or purified mouse laminin isoforms are not commercially available. Nevertheless, given that laminin expression is tightly regulated during different stages of development as well as different regions of the brain, our findings with laminin isoforms support our hypothesis that regulation of local protein synthesis is dependent on ECM composition.

Previous research from our lab suggests that adhesions within growth cones serve as platforms for local translation (Kershner and Welshhans, 2017). Within growth cones, these adhesions link the extracellular matrix to the intracellular actin cytoskeleton, thereby mediating the force needed for growth. Thus, we hypothesize that laminin may bind to and signal through integrin receptors, which are key components of growth cone adhesions. Integrin receptors also have distinct isoforms, and their expression and distribution are also tightly regulated in development and adulthood (Yao, 2025, Nirwane and Yao, 2019). Other widely known non-integrin laminin receptors include dystroglycans, which are equally important for laminin-mediated signaling and are developmentally regulated, and found in growth cones (Yao, 2025, Lindenmaier et al., 2019). Thus, rather than acting merely as a growth-promoting substrate, different laminin isoforms may activate or repress translation, depending on the signaling pathways they trigger through their respective receptors. Our current study suggests that laminin and BDNF signaling not only affects local translation, but also the number of adhesions (Fig. 4D). Future research will identify the receptors and signaling pathways through which laminin activates or represses local translation, and the role of adhesions in this process.

It is important to note that although we found increased generalized translation in growth cones in response to both low-concentration substrate laminin and high-concentration soluble laminin (Fig. 1F and 4E), this does not necessarily mean that there is an increase in the same subset of locally translated mRNAs. As proof of this, we examined the well-studied *β-actin* mRNA, whose local synthesis is particularly important for cytoskeletal reorganization and growth cone turning during axon guidance (Leung et al., 2006a, Yao et al., 2006). Interestingly, low concentrations of laminin, when presented as substrate, increased β-actin local synthesis in the presence of BDNF (Fig. 3). Surprisingly, BDNF alone was not enough to cause this increase. Moreover, a high concentration of laminin applied acutely with BDNF led to a significant decrease in β-actin local synthesis (Fig. 5) despite increasing total translation in growth cones (Fig. 4C and 4E). These opposing effects suggest that laminin does not merely increase or decrease translation but may fine-tune the translation of different mRNAs. This could be mediated through a concentration-dependent activation of differential signaling pathways, or a difference in receptor activation.

Previous studies have demonstrated that laminin distribution is spatiotemporally regulated and that it has a role in axon guidance (Miner et al., 1997, Lentz et al., 1997, Hopker et al., 1999, Lathia et al., 2007, Clark et al., 1993). Laminin distribution *in vivo* during development is patchy (Zhou, 1990, Liesi, 1985, Liesi and Risteli, 1989, Luckenbill-Edds, 1997); therefore, we added laminin via bath application to mimic this pattern in a growth cone encountering laminin. Previous studies have also demonstrated that the concentration of laminin 111 can alter the response of *Xenopus* growth cones to netrin-1, converting attraction to repulsion (Hopker et al., 1999). Because we found that BDNF plus low concentrations of laminin leads to local translation of *β-actin* mRNA (Fig. 3D), but high concentrations did not (Fig. 5C), our studies suggest that local translation may also contribute to this change in growth cone responsiveness.

Interestingly, the amount of *β-actin* mRNA local translation in growth cones is thought to be small compared to the pre-existing β-actin protein monomeric pool (Strohl et al., 2017, Turner-Bridger et al., 2018); yet, local translation is necessary for axon guidance in response to BDNF and netrin-1 (Yao et al., 2006, Leung et al., 2006a, Welshhans and Bassell, 2011). This raises the question of why newly translated β-actin is functionally needed for axon guidance. Our data shows that laminin is necessary for local translation of *β-actin* mRNA, and laminin signaling has mainly been shown to act through adhesions in growth cones; taken together, this suggests that local translation of β-actin is functionally needed at growth cone adhesions to either help create new adhesions or strengthen existing adhesions and thus provide the force needed to mediate axon turning. However, additional studies, both examining this functionally and elucidating the specific signaling pathway, are needed to validate this hypothesis. Taken together, this research demonstrates that laminin and BDNF work through synergistic signaling pathways to induce local translation in developing axonal growth cones. Furthermore, this study demonstrates that laminin isoforms and concentrations may play a key role in directing the amount, location, and specific mRNAs that are translated. Future work will elucidate how local translation of specific mRNA populations is activated or repressed by differential signaling involving the extracellular matrix.

## MATERIALS AND METHODS

### Cell culture

All experimental procedures were approved by the Institutional Animal Care and Use Committee at the University of South Carolina. C57BL/6J male and female mice purchased from the Jackson laboratory (000664) were bred to obtain embryonic day 17 (E17) embryos from timed pregnant females. Dissection was performed to isolate cortices from E17 embryos of either sex. Next, based on the experimental design, cortical tissue was triturated and plated in MEM (Thermo Fisher, 45000-380) with 10% FBS (Sigma, F2442) or Neurobasal media (Thermo Fisher, 21103-049) containing GlutaMax (Thermo Fisher, 35050061) and B27 (Thermo Fisher, 17504001). Neurons were plated on either 100 μg/mL Poly-l-lysine (PLL; Sigma-Aldrich, P1274) or PLL plus 10 μg/mL laminin-coated (Thermo Fisher, 23017015) glass coverslips (Carolina Biological, 633031). Depending on the experiment, either acid-rinsed coverslips, pre-made live cell dishes (MatTek, P35G-1.5-14-C), or lab-made live cell dishes (Corning, 25382-064) were used. For experiments using soluble laminin stimulation, neurons were cultured on 100 μg/mL PLL alone. In all cases, after 2 h of plating, cells were checked for adherence, and the plating media was replaced with Neurobasal with Glutamax and B27. Cells were cultured for 2 days in vitro (DIV) at 37°C in 5% CO_2_ until most neurons had long neurites and an identifiable axon, which was defined as three times as long as the next longest neurite.

### Starve and stimulate paradigm and puromycin assay

To assess total protein synthesis in growth cones, we performed a puromycin assay. After E17 cortical neurons were cultured for 2 DIV, they were starved for 3 hours in Neurobasal media containing Glutamax but no B27. At 2 hours and 30 min into starvation, the negative control group received 40 μM anisomycin (Sigma, A9789) for the remainder of the starvation period. Anisomycin was also included during the stimulation period (40 - 60 min, depending on the experiment). For soluble laminin stimulation experiments, at 3 hours, starvation media was removed, and cells were stimulated with Neurobasal media containing Glutamax and B27, and vehicle (50 mM Tris-HCL (pH 7.4), 0.15 M NaCl) or 25 μg/mL laminin 111 for 15 min, followed by 100 ng/mL BDNF (Thermo Fisher, 450-02) or vehicle for BDNF [0.1% Bovine Serum Albumin (BSA)] for 10 min (Figure 4A and 5A). For experiments with neurons growing on PLL + laminin-coated coverslips or live cell dishes, cells were stimulated with Neurobasal media containing Glutamax and B27, and 100 ng/mL BDNF or vehicle for BDNF (i.e., 0.1% BSA), for 10 min (Figure 1A). Following stimulation, cells were treated with 4 μM puromycin (Millipore Sigma, P8833) or 5 μg/mL O-propargyl puromycin (OPP; ClickChemistry Tools, 1407-25) for 10 min (Figure 1A and 3A) or 30 min (Figure 6A), respectively, depending on the experimental timeline. Cells were washed with PBS and fixed immediately with 4% PFA (VWR, 100504-858) containing 5 mM MgCl_2_ and 4% sucrose for 17 min. Next, cells were washed with PBS containing 2 mM MgCl_2,_ and immunofluorescence was performed. For experiments using OPP, an alkyne-azide click-iT reaction was performed using a click chemistry kit (Vector Labs, CCT*-*1496) with AZDye 647 azide plus as per the manufacturer’s protocol (Vector labs). Timelines for each experiment can be found within its respective figure.

### Laminin isoforms

To examine the effect of various laminin isoforms on local translation in growth cones of E17 mouse cortical neurons, we used human recombinant laminin isoforms 111, 511, 211, and 221 from Biolamina (LNKT-0201). Briefly, neurons were cultured on coverslips coated with 100 μg/mL PLL or PLL plus 10 μg/mL of one of these laminin isoforms. We then performed the starve and stimulate paradigm as described above with slight modifications (Figure 2A). Briefly, after starvation, cells were stimulated with Neurobasal media containing Glutamax and B27 and 100 ng/mL BDNF for 10 min, followed by puromycin treatment for 10 min, and then fixation as previously described.

### Immunofluorescence

Immunofluorescence was performed on all fixed neurons as per a previously published protocol(Welshhans and Bassell, 2011). The following primary antibodies were used: mouse anti-puromycin Alexa fluor 647 conjugate (1:250; Sigma, MABE343-AF647), rabbit anti-paxillin (1:500; Abcam, Ab32084), and rat anti-integrin-beta-1 (1:500; Millipore Sigma, MAB1997). The following secondary antibodies were used: goat anti-rabbit Alexa Fluor 488 (1:1000; Thermo Fisher, A-11034) and donkey anti-rat Cy3 (1:500; Jackson Immuno, 712-165-153). Following secondary antibody washes, coverslips or live cell dishes were briefly washed with 1X PBS and deionized water. Coverslips were mounted with Prolong gold antifade mounting media (Invitrogen, P36934) on slides (VWR, 48311-951). Live cell dishes were stored in fresh PBS until imaging.

### Puromycilation-Proximity Ligation Assay (Puro-PLA)

To examine the local translation of *β-actin* mRNA in cortical neuronal growth cones, we performed PLA in live cell dishes as per the manufacturer’s protocol (Sigma-Aldrich) with slight modifications. Briefly, after labeling nascent peptides with puromycin as mentioned in the puromycin assay above, cells were fixed in 4% PFA-sucrose containing 5 mM MgCl_2_, washed, permeabilized with 0.1%Triton X-100 in PBS for 10 min, and blocked with 4% goat serum in PBS for 30 min. Next, neurons were incubated overnight with primary antibodies to puromycin (1:500 mouse; Kerafast, EQ0001) and β-actin (1:1000 rabbit; Abcam, Ab115777) at 4°C. After three x 5 min washes with PBS, neurons were incubated with rabbit PLA ^plus^ and mouse PLA ^minus^ probes at a dilution of 1:10 in 4% goat serum. Three x 5 min buffer A (0.01 M Tris, 0.15 M NaCl, 0.05% Tween 20) washes were performed, followed by ligation for 30 min at 37°C. Following ligation, three x 5 min buffer A washes were carried out, and the amplification step was performed for 100 min at 37°C. Finally, amplification was stopped using wash buffer B (0.2 M Tris, 0.1 M NaCl pH 7.5) three times for 10 min each. Live cell dishes were stored in buffer B for signal stability at 4°C until imaging.

### Image acquisition and analysis

Imaging of mounted coverslips and live cell dishes for all experiments was performed at 100X oil immersion objective on a Nikon-Ti2 Eclipse microscope with a Hamamatsu ORCA-Fusion camera and Lumencor Sola box. Large images for axon length were taken on the same microscope using a 20X objective. Imaging parameters, such as exposure time and LED intensity, were kept consistent across groups and experimental sets. Fluorescence intensity, puncta number, and axon length were all measured using tools in FIJI software.

For fluorescence intensity of puromycin in growth cones, the polygon tool on FIJI was used to trace the growth cone and measure the mean gray value. Axon length was also measured in FIJI using the line tool. An axon was defined as the longest neurite that is thrice the length of the next longest neurite. To measure axon length, a line was traced along the start of the axon from the cell body to the end of the axon tip or shaft just before the growth cone.

To count the number of paxillin-rich puncta in growth cones, fluorescently labeled paxillin puncta were counted manually. The puncta number was then divided by the growth cone area to get puncta/area values. For Puro-PLA analysis of newly translated *β-actin* mRNA in growth cones, we used the ‘Analyze particle’ tool in FIJI. Before using this tool, the growth cone area was traced using the polygon tool and duplicated. An automated thresholding algorithm called ‘Moments’ was applied to the duplicated image. Next, a mask was created, and under the ‘Analyze’ tool in FIJI, measurements were set to Area, bounding rectangle, limit to threshold, and mean gray value. Finally, under Analyze, ‘analyze particle’ was selected, and ‘display results,’ ‘summarize,’ and ‘add to manager boxes’ were checked and applied. An ROI manager with individual puncta ROI was created. Puncta intensity was measured by selecting each individual ROI and hitting ‘Measure’ present in the ROI manager box. Two puncta in close proximity that could not be separated with the watershed tool were counted as one. Puncta per growth cone for Puro-PLA were measured as described earlier in this paragraph. Mean gray value, axon length, puncta intensity, and puncta per growth cone area for all groups were normalized to the control group and then analyzed and plotted in GraphPad Prism software.

### Statistical analyses

All statistical Analyses were conducted in GraphPad Prism software with signficance set at p β 0.05. Data were assessed for normality in Prism, and the appropriate statistical test was then applied. Statistical tests used in each experiment are described in the figure legend. Error bars in the figures represent the standard error of the mean of three independent experiment sets unless otherwise stated.

## ACKNOWLEDGEMENTS

We thank Jeff Twiss and the Twiss lab for providing helpful comments on this research.

## Competing Interests

No competing interests declared.

## Funding

This work was supported by the National Institutes of Health (R01NS125146 to KW), the Whitehall Foundation (2022-08-023 to KW), and an ASPIRE I grant from the University of South Carolina (to KW).

## Notes

### Competing Interest Statement

The authors have declared no competing interest.

